# Genetic structure and sex-biased gene flow in the history of southern African populations

**DOI:** 10.1101/237297

**Authors:** Vladimir Bajić, Chiara Barbieri, Alexander Hübner, Tom Güldemann, Christfried Naumann, Linda Gerlach, Falko Berthold, Hirosi Nakagawa, Sununguko W. Mpoloka, Lutz Roewer, Josephine Purps, Mark Stoneking, Brigitte Pakendorf

## Abstract

**Objectives:** We investigated the genetic history of southern African populations with a special focus on their paternal history. We reexamined previous claims that the Y-chromosome haplogroup E1b1b was brought to southern Africa by pastoralists from eastern Africa, and investigated patterns of sex-biased gene flow in southern Africa.

**Material and Methods:** We analyzed previously published complete mtDNA genome sequences and ~900 kb of NRY sequences from 23 populations from Namibia, Botswana and Zambia, as well as haplogroup frequencies from a large sample of southern African populations and 23 newly genotyped Y-linked STR loci for samples assigned to haplogroup E1b1b.

**Results:** Our results support an eastern African origin for Y-chromosome haplogroup E1b1b; however, its current distribution in southern Africa is not strongly associated with pastoralism, suggesting a more complex origin for pastoralism in this region. We confirm that the Bantu expansion had a notable genetic impact in southern Africa, and that in this region it was probably a rapid, male-dominated expansion. Furthermore, we find a significant increase in the intensity of sex-biased gene flow from north to south, which may reflect changes in the social dynamics between Khoisan and Bantu groups over time.

**Conclusions:** Our study shows that the population history of southern Africa has been very complex, with different immigrating groups mixing to different degrees with the autochthonous populations. The Bantu expansion led to heavily sex-biased admixture as a result of interactions between Khoisan females and Bantu males, with a geographic gradient which may reflect changes in the social dynamics between Khoisan and Bantu groups over time.

## INTRODUCTION

The extensive genetic, linguistic, and cultural diversity of southern African populations (Barnard, 1992; Cruciani et al., 2002; Wood et al., 2005; Tishkoff et al., 2007; Güldemann, 2008, 2014; Quintana-Murci et al., 2010; Schlebusch et al., 2011) reflects a long history of population movements and interactions. The so-called Khoisan populations are the descendants of some of the earliest humans inhabiting the region; they are foragers and pastoralists who speak indigenous non-Bantu languages characterized by the heavy use of click consonants. We use the term “Khoisan” without any assumption about their genetic or linguistic unity (cf. Barnard, 1992). Three language families are recognized among Khoisan (Figure S1): Tuu and Kx’a, spoken by populations known to have practiced hunting and gathering until recently, and Khoe-Kwadi, spoken by a large number of different ethnolinguistic groups practicing diverse subsistence strategies (Güldemann, 2004, 2005, 2008, 2014; Güldemann and Elderkin, 2010; Heine and Honken, 2010). In addition to the diversity in languages and subsistence strategies, there is broadly-defined phenotypic variation that occurs among Khoisan populations. The majority of individuals speaking Tuu and Kx’a languages, and some of the populations speaking Khoe-Kwadi languages, have light skin pigmentation and relatively short stature (a phenotype we here refer to as the “Khoisan phenotype”). At the same time, the majority of individuals from other Khoe-Kwadi-speaking populations, such as the Damara from Namibia as well as populations inhabiting the Okavango delta and eastern Kalahari, are of taller stature and have darker skin pigmentation (Weiner et al., 1964; Jenkins, 1986). Genetic data revealed that the Khoisan populations harbor some of the earliest branching mtDNA and NRY lineages (Tishkoff et al., 2007; Batini et al., 2011; Rosa and Brehm, 2011; Barbieri et al., 2013b, 2014, 2016). Additionally, autosomal genetic data indicate complex patterns of ancestry for most Khoisan groups, reflecting substantial admixture with other groups as well as between different Khoisan groups (Pickrell et al., 2012, 2014; Uren et al., 2016; Montinaro et al., 2017).

It has been shown that there are at least two admixture events in the history of Khoisan populations that could have contributed to their current genetic ancestry. The earlier admixture event involves a migration from eastern Africa that occurred 900-1,800 years ago (Pickrell et al., 2014; Montinaro et al., 2017; Schlebusch et al., 2017) and is supported by several independent lines of evidence from different disciplines. Archeological data support an introduction of pastoralism from eastern to southern Africa (Mitchell, 2002; Pleurdeau et al., 2012), while based on linguistic data it has been hypothesized that Khoe-Kwadi-speaking populations are the descendants of these pastoralist migrants from eastern Africa (Güldemann, 2008), where livestock is present from 4,000 years ago (Deacon and Deacon, 1999; Phillipson, 2005). Genetic evidence of shared ancestry between the Khoe-Kwadi-speaking Nama pastoralists and the ǂKhomani and Karretjie (whose heritage languages belonged at least in part to the Tuu family), and East African groups, specifically the Maasai, was observed in autosomal data (Schlebusch et al., 2012). Recent studies of ancient DNA from skeletal remains from Africa demonstrated that all modern-day Khoisan groups for which there are genetic data have been influenced by 9-22% genetic admixture from East African/Eurasian pastoralist groups (Schlebusch et al., 2017; Skoglund et al., 2017).

Further evidence for a migration from East Africa comes from lactase persistence variants; it was first suggested by Schlebusch et al., (2012) that the lactase persistence genetic variant in the Nama has an East African origin. In the same study, these observations together with autosomal data were interpreted as support for an East African connection to the Nama. Subsequent studies on lactase persistence confirmed these suggestions by finding elevated frequencies of an East African lactase persistence allele in southern African pastoralist groups and in Khoe-speaking groups, particularly the Nama (Breton et al., 2014; Macholdt et al., 2014, 2015).

In contrast to previous studies that concluded that the spread of pastoralism was driven by demic diffusion, new studies of autosomal data argue that cultural diffusion in the absence of significant gene flow might have played an important role in the spread of pastoralism (and possibly Khoe languages) in southern Africa (Uren et al., 2016; Montinaro et al., 2017). Nevertheless, these studies still find evidence for admixture with Eastern Africa or Eurasian sources ~1,160-1,740 years ago in all the Khoisan populations studied (Montinaro et al., 2017). Further evidence comes from mtDNA haplogroup L4b2, found in Nama and ǂKhomani San as well as in high frequency in the Hadza and Sandawe from Tanzania (Knight et al., 2003; Tishkoff et al., 2007), who also make use of click consonants in their languages. In addition to L4b2, mtDNA haplogroups L5 and L3d might represent a relic of the immigration of eastern African pastoralists, but they are found in very few southern African Khoisan populations and their potential link with eastern Africa is not unambiguous (Barbieri et al., 2014); thus they do not provide compelling evidence of a genetic link between the Khoe-speaking populations and eastern Africa.

In contrast to mtDNA, support for a demic diffusion from East Africa is more obvious when the Y chromosome is considered. Based on the distribution of Y chromosome haplogroup E1b1b (E-M293) and associated microsatellite diversity, it has been suggested that this haplogroup spread through Tanzania to southern-central Africa independently of the migration of Bantu-speaking peoples along a similar route, with a movement of people who brought pastoralism ~2,000 years ago (Henn et al., 2008). However, this study was based on a data set that included just three populations from southern Africa: the foraging !Xuun, the Khwe who have various modes of subsistence, and agriculturalist South African Bantu, and thus it was not possible to demonstrate that this haplogroup occurs in higher frequency in southern African pastoralist populations than in forager populations.

The more recent admixture event reconstructed with genomic data is a consequence of the Bantu expansion that started around 5,000 years ago from the current Cameroon-Nigeria border. This expansion is one of the most influential demographic events on the African continent (Grollemund et al., 2015), and led to a sex-biased pattern of admixture between Bantu speakers and the local groups already present in territories settled by Bantu-speaking groups, including forager populations such as Pygmies in Central Africa and the Khoisan groups of southern Africa (Cruciani et al., 2002; Destro-Bisol et al., 2004; Wood et al., 2005; Tishkoff et al., 2007; Verdu et al., 2009, 2013; Quintana-Murci et al., 2010; Schlebusch et al., 2011; Pickrell et al., 2012, 2014). This sex-biased pattern is the result of mating practices that typically involve Bantu males and autochthonous females, but hardly ever involve autochthonous males and Bantu females (Destro-Bisol et al., 2004). This results in the flow of Bantu Y chromosomes into autochthonous communities, or of autochthonous mtDNAs into Bantu communities, depending on where the children were raised.

Several models attempted to explain the pattern of genetic diversity seen in current forager populations and surrounding Bantu populations. The Static and Moving Frontiers model (Alexander, 1977, 1984) was based on archeological data, and was later modified and expanded by Marks et al., (2014) using genetic data from southern Africa. This model postulates different degrees of interaction among incoming agro-pastoralist and resident foraging groups in the presence of “static” and “moving” frontiers. The expansion of farming societies and/or their technologies into new regions in the absence of ecological and geographical restrictions is termed the moving frontier phase, while a static frontier phase emerges once the expanding society faces a natural or ecological boundary that prevents further expansion. This model implies limited assimilation of foragers into farming communities during the moving frontier phase of the dispersal process, with increasing likelihood of gene flow with foragers in the static frontier phase. Another model, proposed by Destro-Bisol et al., (2004) attempts to explain the observed pattern of genetic structure in Central African forager Pygmy populations and neighboring agro-pastoralist populations. Similarly to the Static and Moving Frontiers model, their model assumes the occupation of territories inhabited by foragers by agro-pastoralists and a retraction of foragers into less favorable areas. In contrast to the Static and Moving Frontiers model, Destro-Bisol et al. attempt to explain the sex-biased pattern of admixture by proposing a temporal development of asymmetrical gene flow between foragers and agro-pastoralists. According to this model, early phases of contact were characterized by symmetrical gene flow between expanding and local populations, while sex-biased gene flow developed gradually as a consequence of developing social inequalities between populations. Even though the two models focus on different forager populations (Khoisan vs Pygmies) and employ different genetic and cultural processes in order to explain the current genetic makeup of populations, both models address the same phenomenon, and thus could be used as a framework for studying the Bantu expansion into areas previously inhabited by foragers.

In this study we use previously published mtDNA and NRY sequences collected from a large and comprehensive sample of Khoisan and Bantu-speaking populations to investigate the genetic history and structure of southern African populations, with a focus on the impact of the east African and Bantu migrations. Based on discrepancies between linguistic and genetic affiliations, we suggest probable language shifts in several populations. Finally, we demonstrate that the intensity of sex-based gene flow varies considerably in southern African populations, and is related to geography.

## MATERIAL AND METHODS

### Samples

We collected data from two available datasets which analyzed the same population samples. Genomic DNA was obtained from saliva samples of Khoisan and Bantu-speaking populations from Botswana, Namibia and Zambia. The Y chromosome dataset consists of ~900kb sequences from the non-recombining region from 547 individuals belonging to 24 different populations (Table S1, Figure 1; Barbieri et al., 2016), and the mtDNA dataset comprises complete mtDNA genome sequences from 680 individuals belonging to 26 different populations (Barbieri et al., 2014). The NRY sequences from the neighboring Khwe-speaking ║Ani and Buga populations were merged together into a combined ║Ani/Buga population due to their low sample sizes; the mtDNA sequences were similarly merged to be directly comparable to the NRY dataset. We refer to the dataset of 23 populations (17 Khoisan and six Bantu) that overlap between the NRY and mtDNA datasets from Namibia, Botswana and Zambia, as the “NBZ dataset” (Table S2). For analyses of autochthonous genetic structure before non-autochthonous haplogroups arrived in the area, we excluded from the NBZ data sets (both mtDNA and the NRY) all Bantu populations, non-autochthonous haplogroups from Khoisan populations, and Khoisan populations with sample sizes less than eight after removal of individuals with non-autochthonous haplogroups. This filtering resulted in 13 overlapping populations between mtDNA and the NRY, and we refer to this dataset as the “AU-NBZ dataset” (Table S2). The analysis of the intensity of the sex-biased gene flow (ISBGF) included additional data from southern African populations for which both mtDNA and NRY haplogroup frequencies were previously published (Coelho et al., 2009; de Filippo et al., 2010; Henn et al., 2011; Schlebusch et al., 2011; Barbieri et al., 2013a; Marks et al., 2014), and we refer to this as the “SA dataset” (Table S3).

**Figure 1.**
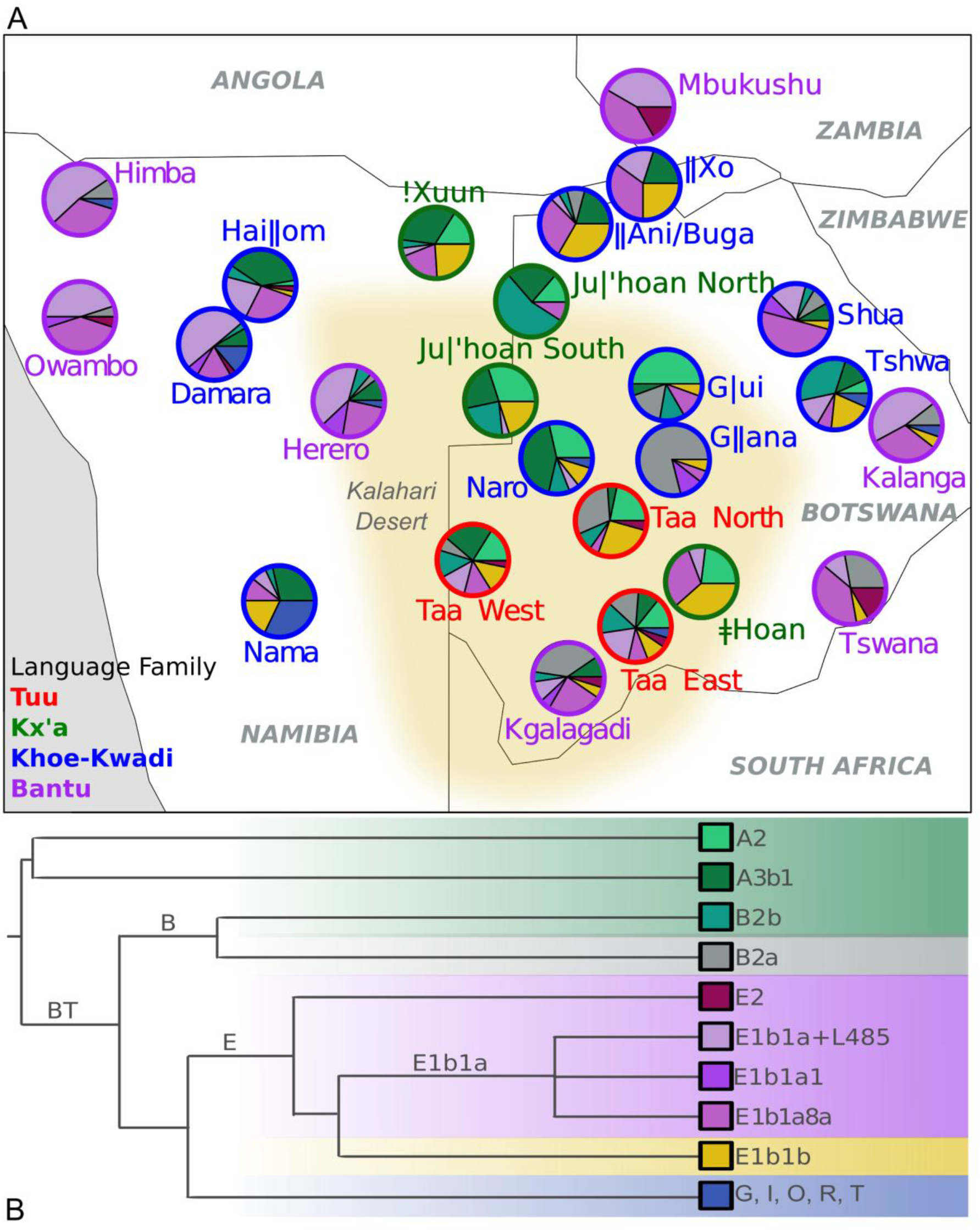
(A) Map of the approximate location of the populations and their NRY haplogroup composition. Population labels are color-coded according to linguistic affiliation as indicated in the lower left corner of the map. Green shades within the pie charts indicate haplogroups that are traditionally considered as Khoisan-related, purple shades indicate Bantu-related haplogroups, yellow indicates E1b1b (thought to be East African), blue are Eurasian haplogroups, and gray represents B2a, which is probably associated with both Khoisan and Bantu populations. The Bantu-related, Eurasian and E1b1b haplogroups are here defined as non-autochthonous haplogroups, while Khoisan-related haplogroups are defined as autochthonous. The yellow area indicates the Kalahari Desert. (B) Schematic representation of NRY phylogenetic tree, from Barbieri et al. 2016.

Since the time of sample collection, additional linguistic research on the Kx’a family has revealed that the language formerly referred to as ǂHoan consists of three dialects: N!aqriaxe, ǂHoan, and Sasi. The language is nowadays referred to as ǂ’Amkoe (Güldemann, 2014; Gerlach, 2016). Although the samples included under the name ǂHoan mainly stem from N!aqriaxe speakers and include only a few ǂHoan speakers, for ease of comparison with previous studies of these samples we continue to refer to them as ǂHoan speakers.

Individuals assigned to haplogroup E1b1b were genotyped for a set of 23 STRs using the PowerPlex^®^ Y23 System (Promega, Mannheim, Germany; Table S4) as described previously (Barbieri et al., 2016). In order to place southern African samples in a broader picture and search for possible connections with eastern Africa, they were subsequently merged with publicly available STR datasets for the E1b1b haplogroup from Africa (Tishkoff et al., 2007; Henn et al., 2008; Berniell-Lee et al., 2009; Gomes et al., 2010; de Filippo et al., 2011), resulting in the “E1b1b-STR dataset” (Table S4) with a total of 278 individuals with 10 overlapping STRs (DYS19, DYS389I, DYS389II, DYS390, DYS391, DYS392, DYS393, DYS437, DYS438, DYS439).

### Data analysis

We used previous haplogroup assignments for both the mtDNA and the NRY data (Barbieri et al., 2013b, 2016). Networks for the NRY haplogroups were generated previously by Barbieri et al. (2016) and were here colored by population in order to investigate the relationships among populations. Branches in the NRY networks were numbered according to the tree in Figure 1 of Barbieri et al., (2016). Additionally, Network 5.0.0.1. (Fluxus Engineering, http://www.fluxus-engineering.com) was used to visualize the relationships between the STR haplotypes genotyped for haplogroup E1b1b. The STR-based network was constructed by applying the reduced-median method first and then the median-joining method (Bandelt et al., 1999) on the dataset with STRs weighted according to their mutation rate (Heinila, 2012). Networks were subsequently plotted and colored with Network Publisher.

Analyses of Molecular Variance (AMOVA; Excoffier et al., 1992) and matrices of pairwise Φ_ST_ distances were computed in Arlequin ver. 3.11 (Excoffier et al., 2007) for both the NBZ and AU-NBZ dataset. For the NRY calculations we used a Tamura and Nei (TrN) model (Tamura and Nei, 1993) with a gamma value of 155.8, and for the mtDNA a TrN model with a gamma value of 0.26, as suggested by results obtained with JModelTest2 (Darriba et al., 2012). An AMOVA was performed in parallel for both the mtDNA and NRY to investigate previously proposed groupings and to explore possible factors that might influence genetic structure. Detailed information about the AMOVA groupings and populations included in each of the groups can be found in Table S5. Ecotype information was obtained from the WWF Terrestrial Ecoregions of the World (TEOW) map (Olson et al., 2001) and populations were grouped accordingly. Nonmetric multidimensional Scaling (MDS) analyses were performed with the function “isoMDS” from the package MASS (Venables and Ripley, 2002). Neighbor-joining trees (NJ trees) depicting population relationships were generated from the matrix of Φ_ST_ distances with the function “nj” of the package ape (Paradis, 2010). Correspondence analysis (CA) was performed with the package ca (Nenadic and Greenacre, 2007), using haplogroup frequencies. A Mantel test was performed between genetic (Φ_ST_) and geographic distances with the R package ade4 (Chessel et al., 2004); geographic distances between populations were averaged over GPS data from the individual sampling locations with the function rdist.earth of the package fields (Furrer et al., 2012).

The intensity of sex-biased gene flow (ISBGF) was calculated for the SA dataset as the difference between the proportion of autochthonous mtDNA haplogroups (L0d, L0k) and autochthonous NRY haplogroups (A2, A3b1, B2b) for each of the populations after removing Eurasian haplogroups in order to avoid recent sex-biased gene flow from European colonists (i.e. NRY haplogroups G, I, O, R, T): ISBGF = %(L0d+L0k) - %(A2+A3b1+B2b). It is not clear if NRY haplogroup B2a is autochthonous to southern Africa or was brought by the Bantu expansion (or possibly both; see Barbieri et al., 2016); we therefore treated haplogroup B2a as ambiguous and ran analyses in parallel treating it as either autochthonous (AUT) or non-autochthonous (NAUT). Values close to zero indicate equal proportions of autochthonous uniparental markers in a given population; positive values indicate a higher proportion of autochthonous mtDNA haplogroups than autochthonous NRY haplogroups; negative values indicate a higher proportion of autochthonous NRY haplogroups than autochthonous mtDNA haplogroups. The p values for the AMOVA, Mantel Test and correlations between ISBGF and longitude and latitude were corrected for multiple testing with the “fdr” method using the function p.adjust from the R package stats (R Core Team, 2014).

#### Differences in migration rates/effective population sizes between females and males

Assuming an island model with neutrality (Wright, 1951; Wilder et al., 2004) one can use mtDNA and NRY F_ST_ values (or Φ_ST_) to explore differences in migration rates between females and males and/or differences in their effective population sizes. The island model with neutrality predicts for a given genetic system that F_ST_ (or Φ_ST_) =1/(1+2ν), where the effective number of migrants is ν =N*m*, with N the effective number of chromosomes (mtDNA or NRY) and *m* the rate of migration. We use the ratio of Nm estimates for mtDNA vs. the NRY to provide insights into the relative amount of female to male migration (Wilder et al., 2004).

## RESULTS

### Y chromosome lineages in southern Africa

#### Khoisan-related haplogroups

Traditionally, the A2, A3b1 and B2b Y chromosome lineages are described as haplogroups characteristic of the autochthonous southern African Khoisan populations of foragers and pastoralists (Underhill et al., 2000; Wood et al., 2005; Quintana-Murci et al., 2010; Batini et al., 2011; Barbieri et al., 2016). Even though A2 and B2b are found both in rain forest forager “Pygmy” populations of the Central African and southern African Khoisan populations, these haplogroups are represented by Pygmy- and Khoisan-specific subclades (Batini et al., 2011). In our dataset Khoisan-related haplogroups are, as expected, observed in higher frequencies in Khoisan than in Bantu populations (Figure S2). However, these haplogroups are also observed in relatively high frequency (>14%) in two Bantu-speaking populations (Herero and Kgalagadi; Table S6).

The A2 haplogroup has a narrow distribution in the central Kalahari (Figure 1, Table S6), and it has been detected in low frequency in Baka foragers from Cameroon and Gabon (Batini et al., 2011). It is mostly present in Kx’a and Tuu-speaking populations, and to a lesser extent in Khoe-Kwadi populations (Hai║om, G|ui, Tshwa and Naro). In the network of A2 sequences (Figure S3), all but one haplotype of the Tuu-speaking populations occur on branch 9 (see Table S1 of Barbieri et al., (2016) for more information about branches), while Kx’a and Khoe-Kwadi populations are present in most of the other branches. All !Xuun, Ju|’hoan South and Ju|’hoan North haplotypes occur in the same branches as Naro haplotypes (for reasons of simplicity from here on we refer to Ju|’hoan North and Ju|’hoan South together as Ju|’hoan; and to Taa West, Taa North and Taa East together as simply Taa). The ǂHoan haplotypes are more closely related to haplotypes from neighboring Taa and G|ui populations than to haplotypes from other Kx’a speakers.

Haplogroup A3b1 shows strong regional and linguistic clustering. It represents a major autochthonous NRY haplogroup in most Khoe-Kwadi-speaking populations (ranging in frequency from 0-43%; Table S6), and is the only autochthonous haplogroup in the Nama. Interestingly, almost all haplotypes from Khoekhoe speakers (Nama, Hai║om and Damara) and !Xuun are found in a single branch of the network (branch 13; Figure S4). Another branch (branch 11; Figure S4) almost exclusively contains haplotypes found in the Khoe-Kwadi-speaking Khwe (║Xo, and ║Ani/Buga) and Naro, which inhabit the Okavango delta and neighboring Ghanzi District, respectively. The Ju|’hoan and Taa populations make up the majority of the haplotypes found in branch 18, and they harbor similar yet distinct sub-lineages within this branch. The Ju|’hoan sub-lineage of A3b1 also comprises one haplotype from Naro and one from !Xuun. Branches 17 and 15 contain haplotypes belonging to populations from all of the language families, and most of the haplotypes from the Bantu-speaking Herero and Kgalagadi.

Present in almost all Khoisan populations (except ║Xo, G║ana and ǂHoan), B2b has frequencies higher than 23% in Ju|’hoan and Tshwa, and frequencies lower than 15% in the other populations (Table S6). All but two haplotypes from the Kx’a-speaking Ju|’hoan and !Xuun belong to branch 26 in the network (Figure S5), which they share with all Naro haplotypes and one Hai║om haplotype. All Taa haplotypes are found in two distinct sub-lineages within branch 26 (Figure S5).

#### B2a haplogroup

Although the B2a haplogroup was previously presented as an indicator of Bantu gene flow (Beleza et al., 2005; Berniell-Lee et al., 2009; Quintana-Murci et al., 2010; Batini et al., 2011), Barbieri et al. (2016) showed that this haplogroup most probably existed in Khoisan populations before the arrival of Bantu speakers. The highest frequency of this haplogroup (~80%) is found in the G║ana (Figure 1, Table S6). In addition, B2a is also found in three other Khoe-Kwadi populations (║Ani/Buga, G|ui and Shua), all Tuu-speaking populations, and all Bantu populations except Mbukushu. It is absent from all Kx’a-speaking populations and the remaining Khoe-Kwadi populations. The most distinct haplotype, closest to the root, is found in a Taa West individual (Figure S6). All of the haplotypes from the West Bantu-speaking Himba, Herero and Owambo are in a specific sub-lineage separate from other haplotypes (dotted circle in Figure S6), while the East Bantu-speaking Kalanga, Tswana and Kgalagadi haplotypes are found in a star-like cluster together with haplotypes from Khoisan populations (mostly from central Kalahari Taa and G║ana populations).

#### Bantu-related haplogroups

Bantu-associated haplogroups such as E1b1a and E2 (Quintana-Murci et al., 2010; de Filippo et al., 2011; Barbieri et al., 2016) are found at frequencies higher than 66% in all Bantu populations except Kgalagadi (43%), while in Khoisan populations their frequency ranges between 3 and 75%, with an average of 29% (Figure 1, Figure S2, Table S6). The most striking pattern is that most of the Tuu and Kx’a-speaking groups have low proportions of Bantu-related haplogroups (range 3-38%), while Khoe-Kwadi-speaking groups vary much more (range 5-75%).

The network for haplogroup E1b1a+L485 sequences (Figure S7) shows a star-like pattern, suggestive of population expansion, that harbors haplotypes from all language families. The Damara haplotypes are found within branch 38, and they are similar to the haplotypes found in neighboring West Bantu-speaking Himba, Herero and Owambo. In the center of the network are haplotypes from West Bantu-speaking populations (Himba, Herero, Owambo and Mbukushu) and Damara, which all come from the northern part of the studied area. Interestingly, a newly described sub-lineage (Barbieri et al., 2016) of this haplogroup (branch 35) is present exclusively in Khoe-Kwadi-speaking groups, namely Khwe (║Ani/Buga and ║Xo) and Hai║om. The similarity of Damara and West Bantu haplotypes is also noticeable in branch 39 of the haplogroup E1b1a1 network (Figure S7). The network for haplogroup E1b1a8a sequences is similar to that of haplogroup E1b1a+L485 in exhibiting a star-like pattern with haplotypes from all of the language families in the core (Figure S8).

Finally, haplogroup E2 is found in frequencies lower than 5% in Taa, Damara, Hai║om, Owambo, and Kgalagadi, while in Mbukushu and Tswana it reaches almost 17% (Figure 1, Table S6). The network for haplogroup E2 sequences shows a Hai║om haplotype closest to the root, while Taa haplotypes stem from Bantu haplotypes (Figure S9).

#### E1b1b haplogroup

This haplogroup is considered to have an East African origin, and it has been associated with the spread of pastoralism from East Africa to southern Africa (Henn et al., 2008; Trombetta et al., 2015). It is almost absent in Bantu populations in our data set (except Kalanga, Tswana and Kgalagadi, where it ranges between 4.7 and 5.6%; Figure 1, Figure S2, Table S6), while it is present in all Khoisan populations except Damara and Ju|’hoan North (ranging between 2.7% and 38.5%, where present). The sequence-based network of this haplogroup shows a star-like pattern with all language families represented in the core of the network (Figure S10). Interestingly, most of the Khoekhoe-speaking Nama haplotypes are found in the core. Haplotypes found in the ǂHoan individuals are on a branch shared with Taa and G|ui haplotypes (dotted circle in Figure S10). The analysis of this haplogroup based on STR data and its possible link to the spread of pastoralism is discussed in detail below.

#### Eurasian-related haplogroups

In eight populations we found 20 individuals (Table S1) with NRY haplogroups that are traditionally considered to be of Eurasian origin (Underhill and Kivisild, 2007). Haplogroups G2a2b2a (n=3), I1 (n=3), and R1a1 (n=1) are found exclusively in the Nama; I2a2a (n=4) and O (n=1) are found exclusively in the Damara; while G2a2b2b (n=1) is found in the Taa East. Other haplogroups are more widespread, e.g. R1b1 is found in five individuals from different populations (one in each of Himba, Kalanga, Tshwa, Nama, and Naro), while T1a2 is found in two individuals (one Nama and one Herero).

#### AMOVA

Factors that might influence the genetic structure of southern African populations can be explored by grouping populations using various criteria and then examining the proportion of genetic variance shared among groups, among populations within groups, and within populations, using AMOVA (Excoffier et al., 1992). Overall, the genetic differentiation among groups is slightly larger for mtDNA than for the NRY (~21% vs. ~17%; Table 1). When all Khoisan populations are analyzed together as one group in the AMOVA, they show slightly higher differentiation between populations for mtDNA than for the NRY (~17% vs. ~15%; Table 1), which may reflect geographically structured mtDNA lineages and recent expansion of Bantu-related NRY haplogroups. However, this pattern varies when the Khoisan language families are analyzed separately; the Khoe-Kwadi harbor the biggest proportion of between-population variance for both uniparental markers. Overall, Bantu groups harbor levels of between-group variation that are comparable to Khoisan populations for mtDNA but lower for the NRY (Table 1), in keeping with other evidence that the Bantu expansion largely involved males (Cruciani et al., 2002; Destro-Bisol et al., 2004; Wood et al., 2005; Tishkoff et al., 2007; Verdu et al., 2009, 2013; Quintana-Murci et al., 2010; Schlebusch et al., 2011; Pickrell et al., 2012, 2014).

**Table 1.** Analysis of Molecular Variance (AMOVA) for the mtDNA and NRY data. The names of different groupings are followed by the number of groups defined in brackets (see Table S5 for information on populations included in each of the groups). FDR-corrected p-values significant at the 0.05 and 0.01 levels are indicated with (*) and (**), respectively.

| mtDNA |  |  | NBZ AMOVA Grouping | NRY |  |  |
| --- | --- | --- | --- | --- | --- | --- |
| Among Groups | Among Populations within Groups | Within Populations |  | Among Groups | Among Populations within Groups | Within Populations |
|  | 21.26** | 78.74 | All populations (1) |  | 17.08** | 82.92 |
|  | 17.27** | 82.73 | Khoisan (1) |  | 14.69** | 85.31 |
|  | 18.18** | 81.82 | Khoe-Kwadi (1) |  | 18.79** | 81.21 |
|  | 7.35** | 92.65 | Kx'a (1) |  | 10.16** | 89.84 |
|  | 8.12** | 91.88 | Tuu (1) |  | 1.17 | 98.83 |
|  | 16.61** | 83.39 | Bantu (1) |  | 8.29** | 91.71 |
| 20.28** | 8.74** | 70.98 | Phenotype (2) | 13.37** | 9.03** | 77.6 |
| 18.46** | 5.37** | 76.17 | Uren (5) | 12.6** | 6.33** | 81.07 |
| 1.57 | 19.91** | 78.52 | WWF (7) | 1.24 | 16.01** | 82.75 |
| 14.61** | 10.46** | 74.93 | Subsistence (4) | 8.15** | 10.96** | 80.89 |
| 18.86** | 3.9** | 77.24 | Geo-linguistic (8) | 9.4** | 8.49** | 82.11 |
| 13.89** | 14.75** | 71.36 | Linguistics (2) | 9.46** | 12.57** | 77.97 |
| 8.77** | 14.49** | 76.74 | Linguistics (4) | 5.84* | 12.58** | 81.58 |

The highest values of between-group variance (~20% for mtDNA and ~13% for the NRY) is seen when grouping populations by two broadly-defined phenotypes (“Khoisan” and “Non-Khoisan” phenotypes), indicating that this phenotypic distinction reflects different population histories. The second highest between-group variance for the NRY, and one of the highest for mtDNA, is seen when populations are grouped into five groups defined by distinct, geographically organized autosomal ancestry components inferred from unsupervised population structure analysis (Uren et al., 2016). Based on the distribution of autosomal ancestries they defined, Uren et al., (2016) concluded that the autosomal structure in Khoisan populations reflects the role of geographic barriers and the ecology of the greater Kalahari Basin. In order to test if our dataset of uniparental markers is in agreement with this conclusion, we obtained ecotype information from the WWF Terrestrial Ecoregions of the World (TEOW) map (Olson et al., 2001) and grouped populations accordingly (WWF, Table 1). The much lower among-group variance for this grouping (for both uniparental markers) suggests that the ecoregions in which the populations currently live does not explain the genetic structure of the studied populations. Grouping populations by the geo-linguistic categories defined by Barbieri et al., (2014) results in capturing ~19% of the variance seen in mtDNA between eight geo-linguistic categories (Geolinguistic, Table 1) and is thus one of the best groupings for the mtDNA, but it captures just ~9.4% variation for the NRY. When only linguistic criteria are applied, grouping by the two major linguistic groups (Bantu and Khoisan) better explains the variance between groups for both mtDNA and the NRY than when grouping by the four language families (Bantu, Kx’a, Tuu and Khoe-Kwadi).

The results of the AMOVA analysis for the AU-NBZ data set, i.e. after removing the Bantu populations and non-autochthonous uniparental lineages (Table S7) reveals that grouping by phenotype has very low explanatory power, contrary to the entire dataset (Table 1). This suggests that the association with phenotype in the entire dataset is driven by the difference between autochthonous and non-autochthonous haplogroups. There is also larger genetic differentiation among groups for the NRY than for the mtDNA (~23% vs. ~8%) for the AU-NBZ dataset (comprising only autochthonous lineages), in contrast to the full dataset (~17% vs. ~21%), suggesting differences in male vs. female migration between Khoisan and Bantu groups. Sex-biased gene flow is analyzed in more detail below.

#### Φ_ST_ distance-based analysis: MDS and NJ tree

Bantu-speaking populations are separated from Kx’a and Tuu speakers in both the mtDNA and the NRY MDS (Figure 2), while the Khoe-Kwadi populations are spread between them. As expected, this pattern is also noticeable in the mtDNA and the NRY NJ trees (Figure S11 A-B) where Bantu populations tend to be on the opposite side of the tree compared to Kx’a and Tuu-speaking populations. The Damara appear to be closer to Bantu-speaking populations than to the majority of the Khoisan populations for both mtDNA and the NRY, while the G║ana population is a clear outlier in the NRY MDS, with the Kgalagadi and Taa North as the closest populations. Within the Bantu populations, Kgalagadi and Tswana from the central Kalahari are more distant from the rest of the Bantu populations for both mtDNA and the NRY (Figure 2). Interestingly, the Kgalagadi appear deep within Khoisan branches on the NRY NJ tree (also reflected in the MDS). In contrast to the Kgalagadi and Tswana populations, the northwest Namibian Himba and Herero represent the most distant populations from the Tuu and Kx’a populations in the mtDNA-based tree, while Himba, Kalanga and Mbukushu are distant in the NRY-based tree.

**Figure 2.**
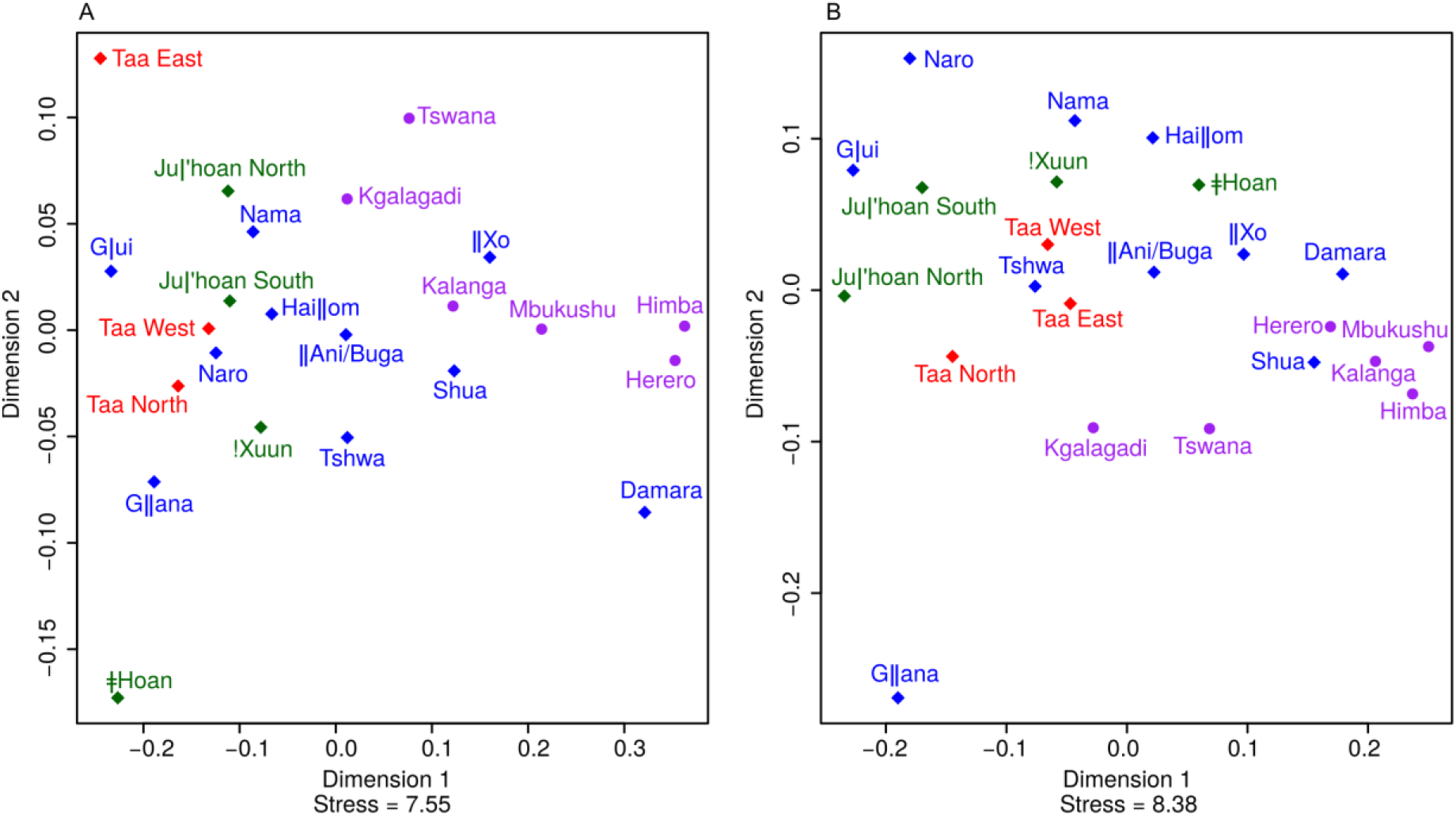
Multidimensional Scaling plot based on mtDNA (A) and NRY (B) Φ_ST_ distances (Table S11). Population symbols and colors indicate linguistic affiliation, as shown in Figure 1.

#### Correspondence analysis (CA)

In contrast to the Φ_ST_ distance-based analyses, the distinction between Khoisan and Bantu populations in the CA (Figure S11 C-D), which is based on haplogroup frequencies, is more striking. Most Khoisan populations exhibit low differentiation for mtDNA haplogroups due to high frequencies of the autochthonous haplogroups L0d and L0k, whereas in the NRY CA plot most of the Bantu populations are clustered together due to their high frequency of Bantu-related NRY haplogroups, such as E1b1a and E2. The Khoe-Kwadi populations are dispersed between Bantu populations on the one side and Kx’a and Tuu populations on the other side. Although the G║ana group with other Khoisan populations in the mtDNA CA plot, they are a clear outlier in the NRY CA plot, due to their high frequency of haplogroup B2a. As in the MDS analysis, the mtDNA and NRY CA plots show the Damara closer to the Bantu-speaking pastoralist Himba and Herero populations than to other Khoisan populations.

#### Mantel Test: geographic vs. genetic distances

There is a statistically significant correlation between pairwise geographic distances and mtDNA Φ_ST_ distances (Mantel Test, R^2^=0.258, FDR-corrected p value=0.02), but not between pairwise geographic distances and NRY Φ_ST_ distances (Mantel Test, R^2^=0.038, FDR-corrected p value=0.32). Together with the AMOVA results (Table 1) showing bigger among-group differences for mtDNA than for the NRY, these results indicate that mtDNA tends to be geographically more localized than the NRY. To exclude the impact of the East African, Bantu and European haplogroups, we performed the Mantel Test between pairwise geographic distances and mtDNA and NRY Φ_ST_ distances from the AU-NBZ dataset. The Mantel Test on this data set still did not show a significant correlation between NRY Φ_ST_ distances and geography (Mantel Test, R^2^=0.266, FDR-corrected p value =0.12), but it did show an increase in R^2^ value. In contrast, mtDNA Φ_ST_ distances and geography showed a moderate decrease in R^2^ value and the correlation was not significant anymore (Mantel Test, R^2^=0.214, FDR-corrected p value =0.14).

#### NRY haplogroup E1b1b and pastoralism

The NRY haplogroup E1b1b has an East African origin and has been associated with the spread of pastoralism from East Africa to southern Africa (Henn et al., 2008). However, this previous study included just three populations from southern Africa (!Xuun foragers, Khwe that practice various subsistence strategies, and South African Bantu agro-pastoralists), and therefore could not test whether the E1b1b haplogroup may have been brought to southern Africa by a pre-Bantu pastoralist migration. With our larger dataset of southern African Khoisan populations that practice a variety of subsistence strategies, we therefore tested if the E1b1b haplogroup was in higher frequency in the pastoralist Nama than in populations with various subsistence practices and foragers. We calculated the proportion of E1b1b in Khoisan populations separately for the pastoralist Nama, groups practicing various subsistence strategies in recorded history (║Ani/Buga, ║Xo, Shua, Tshwa), and for traditional forager populations (Kx’a and Tuu populations, Naro, Hai║om, G|ui and G║ana; classified according to: Widlock et al. not dated, http://dobes.mpi.nl/projects/akhoe/people/; Cashdan, 1986; Barnard, 1992; Chebanne, 2002; Dieckmann, 2014). Even though our results indicate that the pastoralist Nama have the highest frequency of E1b1b, there are no significant differences in frequencies between different Khoisan subsistence groups (Table S8; 3-sample test for equality of proportions without continuity correction p-value = 0.77). In order to exclude possible masking of the pre-Bantu structure we compared frequencies of E1b1b in groups practicing different subsistence strategies after excluding Eurasian and Bantu-related haplogroups (i.e. E1b1a, E2, G, I, K and R1) and Khoisan populations with high proportions of non-autochthonous ancestry (i.e. populations with predominant non-autochthonous uniparental ancestry regardless of treatment of B2a: ║Xo, Shua, and Damara; discussed in more detail in the section “Dominant uniparental ancestry components”). After this filtering the difference between subsistence groups is statistically significant (Table S9; 3-sample test for equality of proportions without continuity correction p-value = 0.0058). Populations with various subsistence strategies have significantly higher frequencies of E1b1b than foragers (2-sample test for equality of proportions with continuity correction FDR-corrected p-value = 0.023). However, the pastoralist Nama do not have significantly different frequencies than foragers or populations practicing various subsistence strategies (2-sample test for equality of proportions with continuity correction FDR-corrected p-values: 0.32; 1, respectively). It should be noted, however, that the sample size for pastoralists in our dataset is low, and overall there is large variation in sample size for each of the subsistence-based groups as well as in the number of populations included in each of the groups. We also tested if the E1b1b haplogroup was in higher frequency in the pooled Khoe-Kwadi populations than in the pooled Tuu or Kx’a populations; the difference in frequency between different Khoisan linguistic groups is not significant (Table S8; 3-sample test for equality of proportions without continuity correction p-value = 0.16). Overall, our data thus do not provide compelling support for a link between haplogroup E1b1b and current pastoralists or Khoe-Kwadi speakers.

#### E1b1b STRs

To further investigate the relationships of southern and eastern African E1b1b Y chromosomes, we genotyped our E1b1b samples for 23 STR loci and merged these with previously-published data (Tishkoff et al., 2007; Henn et al., 2008; Berniell-Lee et al., 2009; Gomes et al., 2010; de Filippo et al., 2011). A network based on the 10 STR loci in common between the studies shows that individuals from East Africa have the highest diversity of haplotypes in the data set, while among the southern African Khoisan populations, Khoe and Kx’a speakers harbor the highest diversity of haplotypes (Figure 3A). The two eastern African foraging populations that speak languages with click consonants, the Hadza and Sandawe, are spread across the network, and they show sharing of haplotypes with southern African populations, suggesting recent gene flow or a common origin of haplotypes in these populations. Besides the Hadza and Sandawe, some haplotypes from Nilotic, southern African Bantu, and (to a lesser extent) Afroasiatic populations are similar to haplotypes found in the Khoisan populations. Haplotypes found in Khoe populations are shared with other Khoisan groups, East African Hadza, Sandawe and Nilotic populations, and southern African Bantu, and are generally either shared or in close proximity to Sandawe haplotypes. The majority (~75%) of the new southern African haplotypes are defined by 10 or fewer repeats at the DYS389I locus. All of the Khoe-speaking Nama individuals carry haplotypes with the DYS389I-10 allele, and three out of four Nama haplotypes are shared either with other Khoe populations, Tuu, southern African Bantu, Sandawe, or Kx’a populations. The segregation between the two clusters in the sequence-based network of haplogroup E1b1b indicates that the DYS389I haplotype with 10 repeats is most likely ancestral (Figure 3B), contrary to previous suggestions (Henn et al., 2008). Overall, the diversity of haplotypes seen in different Khoisan populations, with multiple star-like expansions from haplotypes in close proximity to eastern African foragers, suggests a more complex history for haplogroup E1b1b than previously suspected.

**Figure 3.**
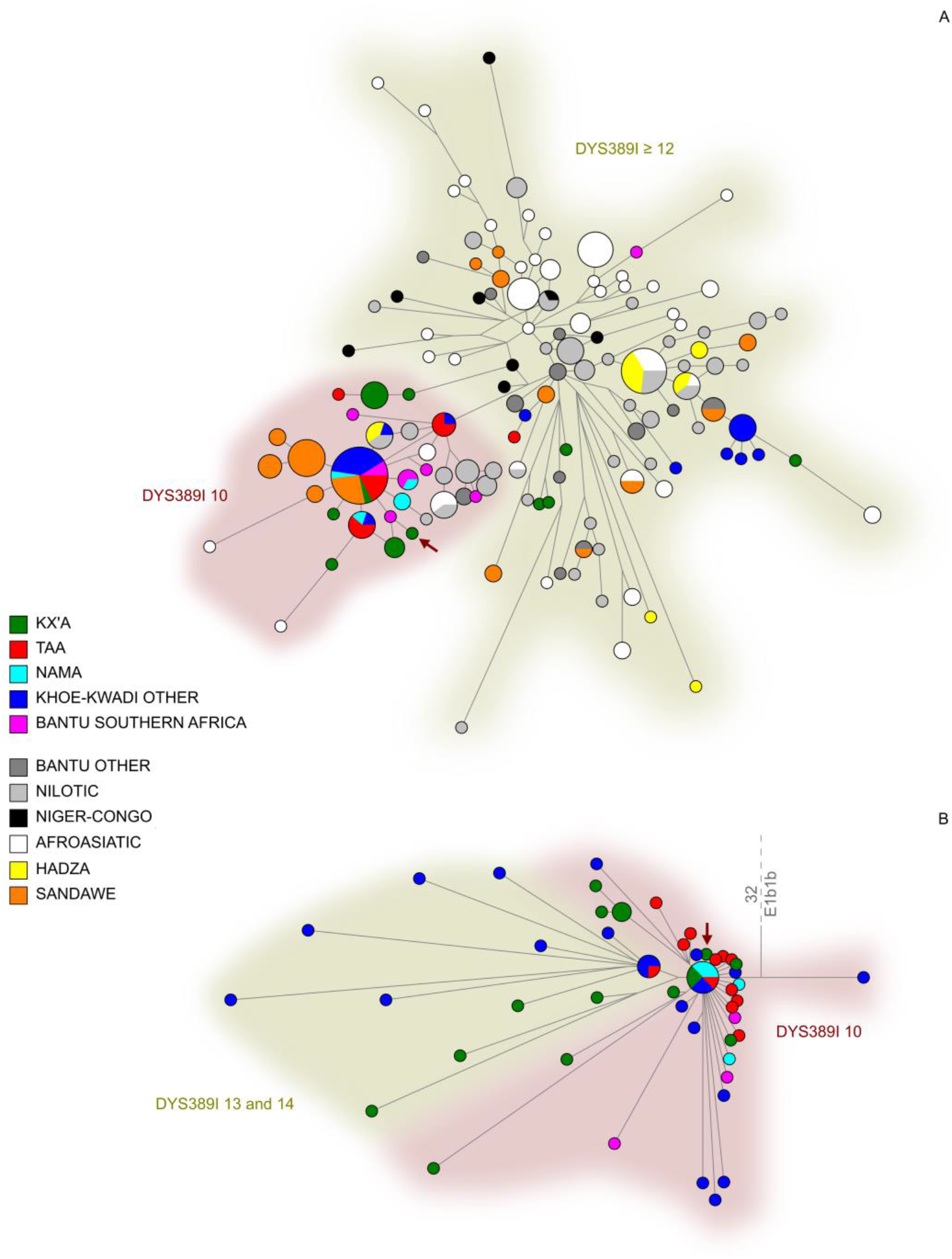
Median-joining networks of (A) 10-locus Y-STR haplotypes for individuals with the NRY E1b1b haplogroup; (B) haplogroup E1b1b sequences (zoomed in from Figure S12 of Barbieri et al. 2016). Shaded areas indicate the number of DYS389I repeats present in the STR haplotypes. The red arrow indicates a haplotype with 9 DYS389I repeats.

### Dominant Uniparental Ancestry Components and the Intensity of the Sex-Biased Gene Flow

#### Dominant uniparental ancestry components

Most of the Khoisan populations are characterised by high frequencies of autochthonous mtDNA lineages and somewhat lower (but still relatively high) frequencies of autochthonous NRY haplogroups (Figure 4, Table S6). However, a different pattern is observed for Kx’a and Tuu populations versus Khoe-Kwadi-speaking populations. All Kx’a and Tuu-speaking populations are characterized by more than 80% autochthonous mtDNA lineages and variable frequencies of autochthonous NRY haplogroups (ranging from 23% to 91% regardless of whether B2a is treated as autochthonous or non-autochthonous), while Khoe-Kwadi populations are characterized by variability in the proportion of autochthonous mtDNA (ranging from 13% to 100%) and NRY haplogroups (B2a as AUT: 11-83%; B2a as NAUT: 0-81%), and thus are more widely dispersed across the plot (Figure 4). Bantu populations are characterized by an excess of autochthonous mtDNA haplogroups, and hence they show female-biased gene flow from autochthonous populations, while in the Khoisan populations the sex-biased gene flow is characterized by an excess of the non-autochthonous Y chromosome haplogroups, showing male-biased gene flow from non-autochthonous populations.

**Figure 4.**
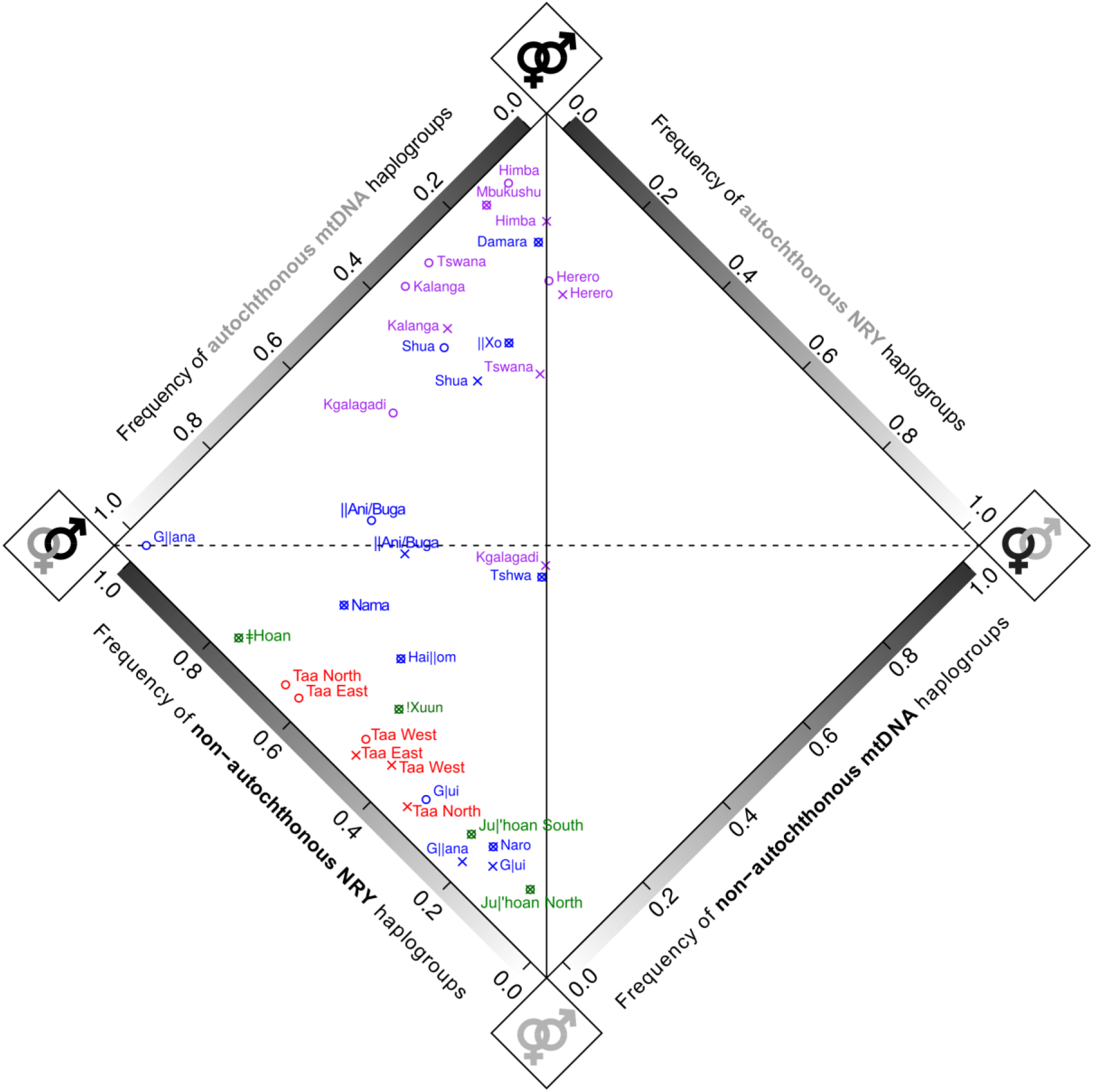
Diamond plot showing dominant uniparental ancestry components based on mtDNA and NRY haplogroup frequencies. Haplogroups of Eurasian origin are included here as non-autochthonous haplogroups, in order to depict all contributions of non-autochthonous origin to each population, but are removed from calculations of the intensity of sex-biased gene flow (see text for further details). Language family affiliation is indicated by the color of the population name (green – Kx’a, red – Tuu, blue – Khoe-Kwadi, and purple - Bantu). The horizontal dotted black line separates populations with predominantly autochthonous haplogroups (below the line) from populations with predominantly non-autochthonous haplogroups (above the line), based on both mtDNA and NRY haplogroup composition. The vertical black line represents equal proportions of non-autochthonous mtDNA and NRY haplogroups (and hence no sex bias), while the distance from this line reflects the intensity of sex-biased gene flow. The B2a haplogroup was treated separately as either autochthonous (“x”) or as non-autochthonous (“o”).

#### Intensity of the Sex-Biased Gene Flow (ISBGF)

Most of the Bantu populations have moderate or no sex-biased gene flow (with a mean of 0.137), while different Khoisan populations show varying degrees of ISBGF (with a mean of 0.386; Figure 5 A and C). The Khoisan populations show significantly higher values for ISBGF than Bantu populations, regardless of the treatment of B2a as autochthonous or non-autochthonous (Wilcoxon rank sum test with continuity correction, B2a as AUT: W = 413, FDR-corrected p value = 0.0007, 95% CI = 0.108-0.354, Figure S12C; B2a as NAUT: W = 387, FDR-corrected p value = 0.003, 95% CI = 0.075-0.377; Figure 5C). Treatment of B2a as autochthonous or non-autochthonous does not strongly influence ISBGF (Figure S12) due to a strong correlation between ISBGF when B2a is treated as autochthonous and when it is treated as non-autochthonous (p=2.6e-13, R^2^= 0.71; Figure S13). The treatment of B2a has the strongest influence on the G║ana, who exhibit the strongest sex-biased gene flow in the SA dataset (ISBGF = 1) when B2a is treated as a non-autochthonous haplogroup, but have a value of 0.21 when it is treated as an autochthonous lineage. Moreover, populations in the central Kalahari and South Africa exhibit stronger ISBGF (Figure 5A), and there is a significant correlation (FDR-corrected p value=1.4e-05, R^2^= 0.31) between latitude (north to south) and ISBGF, but not between longitude and ISBGF (Figure S14). The correlation between ISBGF and latitude remains significant when Khoisan and Bantu populations are analyzed separately (Bantu: B2a as NAUT: FDR-corrected p value=2.04e-05, R^2^= 0.6; B2a as AUT: FDR-corrected p value=0.0034, R^2^= 0.34; Khoisan: B2a as NAUT: FDR-corrected p value=0.0108, R^2^= 0.33; B2a as AUT: FDR-corrected p value=0.0015, R^2^= 0.34; Figure S15). The stronger ISBGF seen in southern populations is driven mostly by higher levels of Khoisan-specific mtDNA haplogroups in southern Bantu populations, and higher levels of Bantu-specific NRY haplogroups in Khoisan populations (Figure 5, Table S6). In Bantu populations there is a statistically significant increase of Khoisan mtDNA lineages from north to south (FDR-corrected p value=7.5e-07, R^2^= 0.69; Figure S16).

**Figure 5.**
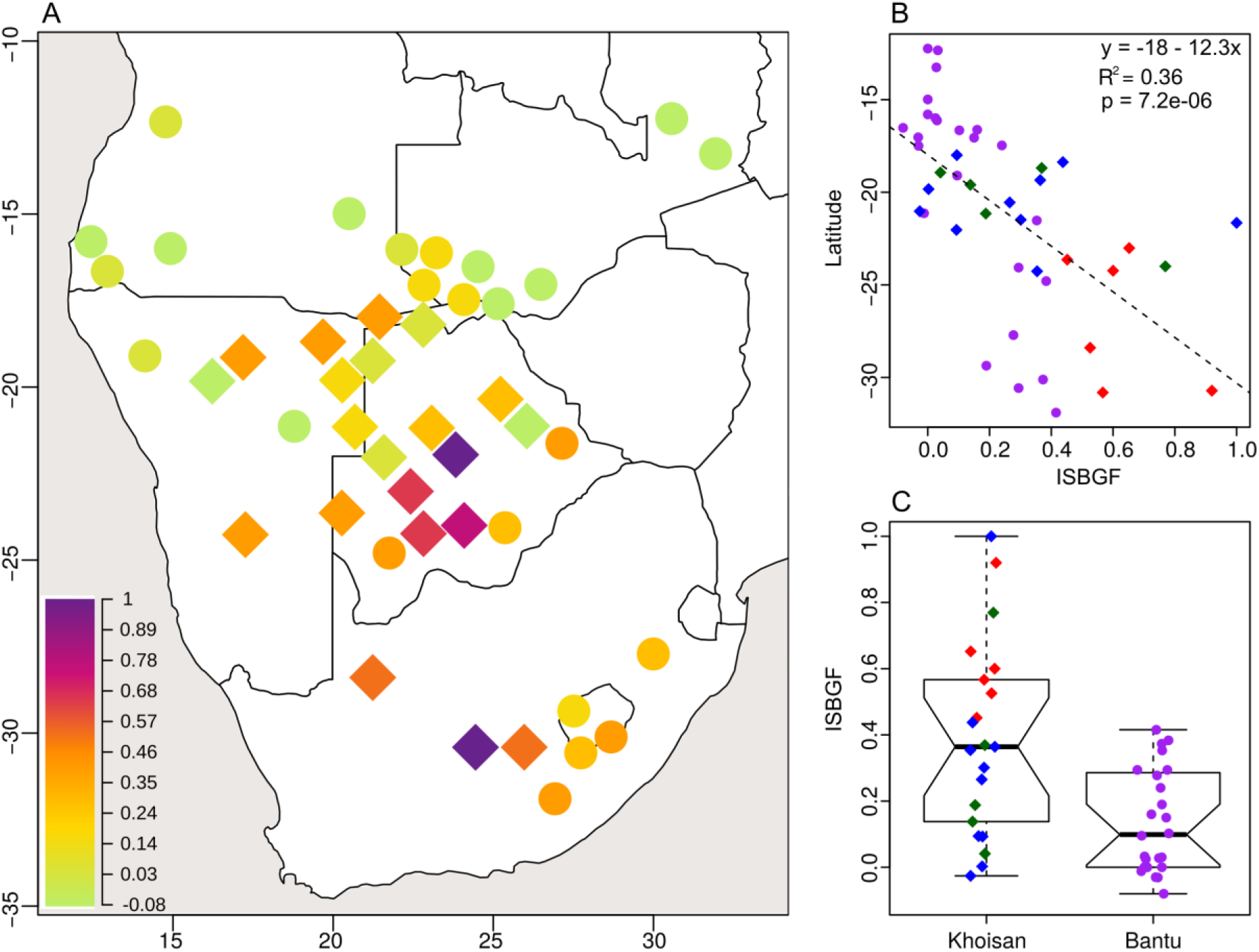
(A) Map of the intensity of sex-biased gene flow (ISBGF) statistic, treating haplogroup B2a as a non-autochthonous haplogroup. Dots and diamonds indicate the approximate geographic location of Bantu and Khoisan populations, respectively. The color indicates the ISBGF value: values close to zero indicate equal proportions of autochthonous uniparental markers in a given population; positive values indicate a higher proportion of autochthonous mtDNA haplogroups than autochthonous Y chromosome haplogroups, and thus male-biased gene flow from non-autochthonous populations to autochthonous populations and female-biased gene flow from autochthonous populations to non-autochthonous populations; negative values (observed only in the Herero) indicate the opposite. (B) Correlation between latitude and ISBGF; C) Differences in ISBGF between Khoisan and Bantu populations (Wilcoxon rank sum test with continuity correction indicates significant differences in the ISBGF means between Khoisan and Bantu populations: W = 387, FDR-corrected p value = 0.003, 95% CI = 0.075-0.377). Different Khoisan language families are represented with different colors as in Figure 1.

#### Differences in migration rates between females and males

Under an island model with neutrality, the F_ST_ genetic distances can be used to estimate the effective number of migrants between populations (see Material and Methods). The ratio of estimated female to male migration (ν_mtDNA_/ν_NRY_) for the entire dataset is 0.796 (Table S10), indicating that the effective number of male migrants is 1.257 times bigger than that of female migrants. When just Bantu populations are taken into account, the ratio becomes even smaller (ν_mtDNA_/ν_NRY_ = 0.443) suggesting that the effective number of male migrants is 2.255 times higher than that of female migrants. In contrast, Khoisan populations show relatively equal migration rates for the mtDNA and NRY, although the ratio varies greatly among the different language families. The overall effective migration rate between Bantu and Khoisan populations is 1.35 times higher for males than for females, which together with the stronger ISBGF in Khoisan populations than in Bantu (Figure 5C) suggests that males from Bantu populations are migrating 1.35 times more frequently to Khoisan populations than Khoisan females are migrating to Bantu populations.

## DISCUSSION

### Factors Influencing the Genetic Structure of southern African Populations

Southern African populations are remarkable for their genetic and cultural diversity as reflected by the wide variety of languages and different subsistence strategies. Here, we compare their maternal and paternal genetic structures and discuss the role of putative factors proposed in previous studies as important in shaping the genetic makeup of the area.

The AMOVA analysis indicates that different factors could be responsible for the current genetic structure based on the uniparental markers of southern African populations. A grouping based on ancestry components detected in autosomal data by Uren et al., (2016) is the best in explaining the variation among Khoisan groups based on only autochthonous haplogroups for the NRY and second-best for mtDNA (AU-NBZ dataset, Table S7), and is also the second-best in terms of explaining the variation among groups for both mtDNA and the NRY for the entire dataset (NBZ dataset, Table 1). Uren et al., (2016) based their grouping on fine-scaled autosomal genetic structure, and interpreted this as reflecting the role of geographic barriers and the ecology of the greater Kalahari Basin. In order to test if this interpretation is consistent with our dataset of uniparental markers and real ecotypes, we grouped populations according to terrestrial ecoregions. Our results indicate that ecology (here defined as WWF ecosystem categories in which the populations live) has very low explanatory power for both uniparental markers (WWF in Table 1). This might be an indication that the structure detected by Uren et al. does not actually reflect ecological boundaries, but rather results from a more complex mixture of geographical and historical factors. In addition, the WWF ecosystem categories are based on modern-day data; these might not accurately reflect the prehistoric climate and vegetation patterns and thus might be poor proxies for the potential environmental factors that shaped the genetic variation seen today in southern Africa.

Overall, the AMOVA groupings with the best explanatory power are based on the differences between Bantu and Khoisan populations, either in phenotype, languages or subsistence strategies (Table 1). The highest value of the between group variance for both uniparental markers is seen in grouping by the two phenotypes (“Non-Khoisan” vs. “Khoisan” phenotype; Table 1), which is expected to correlate with genetic structure to the extent that there is a genetic basis for these broadly-defined phenotypes. Interestingly, Khoisan populations with “Non-Khoisan” phenotypes are also reported to carry a high proportion of autosomal Bantu-related ancestry (Pickrell et al., 2014); these same populations carry high frequencies of non-autochthonous uniparental markers (Figure 4), and they are close to the Bantu populations (Figure 2, Figure S11). The drastically different results obtained in the AMOVA for the grouping based on the two phenotypes for the AU-NBZ (Table S7) and NBZ datasets (Table 1) further indicates that admixture with Bantu populations is the main factor separating populations with different phenotypes. Thus, our results suggest that the major factor influencing the structure of both mtDNA and the Y-chromosome in southern Africa is the distinction between the culturally and genetically divergent Khoisan and Bantu ancestries.

### Fine-scaled population structure

Even though the strongest factor influencing the genetic structure of southern African populations is the distinction between Khoisan and Bantu ancestry, there are also several populations that exhibit substantial frequencies of both ancestries. Many of these speak Khoe-Kwadi languages, and they exhibit quite variable frequencies of autochthonous uniparental markers (Figure 4) as well as very different haplogroup compositions (Figure S11 C-D). Some of the Khoe-Kwadi populations are found to share identical or closely related haplotypes with Bantu populations for both mtDNA and the NRY (Figures S7-S9 and S17-S20), further supporting their close relationship and long history of contact with Bantu populations. We also identify several cases of mismatches between linguistic affiliation and genetic makeup, which are suggestive of language shift. We find putative examples not only of expected shifts from Khoisan to Bantu languages due to the cultural dominance of Bantu-speaking populations, but also putative shifts from Bantu to Khoisan languages, as well as language shifts between Khoisan language families; these are discussed in more detail below.

#### Damara, Himba and Herero

The Damara are an enigmatic group of Khoe-Kwadi speakers; in all analyses they appear to be genetically more similar to Bantu groups (in particular, the Himba and Herero) than to other Khoisan groups (e.g., Figure 1, Figure 2, Figure 4, Figure S11). Notably, 75% of their NRY haplogroups are Bantu-related, while only 11% are autochthonous lineages (Figure 1). With respect to mtDNA, the Damara have a high frequency of haplogroup L3d (63%), which further supports their link to the neighboring Himba and Herero, who also have relatively high frequencies of this haplogroup (Barbieri et al., 2014; Oliveira et al., 2017). The uniparental data are thus in agreement with autosomal genome-wide data that group the Damara with Bantu groups, in particular the Himba and Herero (Pickrell et al., 2012, 2014; Uren et al., 2016; Montinaro et al., 2017). Thus, the current genetic makeup of the Damara population appears to reflect shared ancestry with Bantu-speaking populations (such as Himba and Herero), with subsequent language shift from a Bantu to a Khoekhoe language (Oliveira et al., 2017). Other Khoe-Kwadi populations reported to have high autosomal Bantu-related ancestry (e.g. Shua, ║Xo, and to a lesser extent, Tshwa; Pickrell et al., 2014) also tend to be more similar to Bantu populations based on uniparental haplogroup composition and Φ_ST_ distances, and thus Φ_ST_ based analyses (Figure 2, Figure S11, Table S11). This high level of Bantu-related ancestry, reflecting extensive admixture and/or language shift, needs to be taken into account when considering the relationships of these Khoe-Kwadi populations.

#### Hai║om and Nama

The lowest NRY Φ_ST_ distances for the Khoe-Kwadi-speaking Nama and Hai║om are with the Kx’a-speaking !Xuun (Figure 2, Figure S11, Table S11). The low differentiation between these populations is most probably driven by the sharing or close similarity of sequences from Hai║om, Nama, and !Xuun for NRY haplogroup A3b1 (branch 13 in Figure S4). Overall, the low differentiation between Hai║om and !Xuun would lend some credit to the suggestion (e.g., Barnard, 1992: 12) that some !Xuun speakers shifted to the Khoekhoe language spoken by the Hai║om. The low mtDNA Φ_ST_ distances between the Hai║om and the Nama (Table S11, Figure 2, Figure S11) are in agreement with the proximity of the two languages, as both belong to the Khoekhoe sub-branch of the Khoe-Kwadi family. Interestingly, individual sub-lineages of mtDNA haplogroup L3d (arrow in Figure S17) and NRY haplogroup E1b1a+L485 (Figure S7, branch 35) are found exclusively or predominantly in the Hai║om and other Khoe-Kwadi speakers (especially the Khwe populations from the Okavango delta: ║Ani/Buga and ║Xo), and as such they might represent remnants of lineages that were present in the proto-Khoe-Kwadi population, and they might testify to ancient contact with Bantu populations.

Even though closely related to the neighboring Hai║om, the Nama stand out in having the highest frequency of Eurasian NRY haplogroups (Figure 1). With the exception of one B2b sequence, the only autochthonous NRY haplogroup in the Nama is A3b1, and moreover all Nama sequences fall in just one sub-lineage of this haplogroup (branch 13, Figure S4). This, and the genetic proximity based on NRY Φ_ST_ distances between Nama, !Xuun and Hai║om (Table S11, Figure 2) as well as the network analyses (branch 13 in Figure S4), all suggest that Nama probably incorporated autochthonous NRY haplogroups from the populations living in northern Namibia and Botswana. The connection between Nama and some northern Khoisan populations was previously suggested in anthropological and genetic studies (Nurse and Jenkins, 1977; Barnard, 1992; Boonzaier, 1996; Schlebusch, 2010; Pickrell et al., 2012). On the other hand, Nama autosomal (Uren et al., 2016; Montinaro et al., 2017) and mtDNA (Schlebusch et al., 2013) data suggest their close relationship to southern Khoisan groups such as the ǂKhomani and Karretjie. In our data set the second lowest mtDNA Φ_ST_ distances for Nama after the Hai║om is with Taa West followed by Naro and Ju|’hoan (Table S11). It thus appears that the Nama have a complex genetic history, with different patterns in the NRY and mtDNA suggestive of admixture with different Khoisan populations.

#### ǂHoan

Based on NRY Φ_ST_ distances, the ǂHoan, a Kx’a-speaking population, appear more similar to the Khwe-speaking populations (║Ani/Buga and ║Xo) and to the Tuu-speaking populations than to the Kx’a-speaking Ju|’hoan populations (Figure 2, Figure S11). The closer connection of the ǂHoan with the Khwe-speaking populations is most likely driven by haplogroups E1b1b and E1b1a8a, which are in high frequencies in the ǂHoan (Figure 1). However, network analysis shows that NRY sequences found in the ǂHoan are related to sequences found in the Tuu-speaking Taa populations and in the G|ui (Figures S3 and S10). The closer relationship of the ǂHoan to neighboring Khoe-Kwadi and Tuu-speakers than to other geographically more distant Kx’a-speakers is additionally supported by mtDNA Φ_ST_ distances, as they appear more similar to neighboring Khoe-Kwadi-speaking G║ana and Naro, and Tuu-speaking Taa North and Taa West than to other Kx’a speakers (Table S11). Autosomal data (Pickrell et al., 2012, 2014; Uren et al., 2016; Montinaro et al., 2017) further supports the close connection between Taa and ǂHoan. The genetic data thus provide evidence for long-term extensive contacts between these groups, which is in good accordance with the linguistic data (Gerlach, 2016).

#### Naro

The Khoe-Kwadi-speaking Naro exhibit the smallest NRY Φ_ST_ distances with the Kx’a-speaking Ju|’hoan South (Table S11), and they are not genetically differentiated from the Ju|’hoan South and Tuu-speaking Taa West populations in the mtDNA (Table S11). The low differentiation between these populations is most probably driven by the sharing or close similarity of sequences from Naro, Kx’a and Tuu-speakers in the NRY (Figure S3, branch 26 in Figure S5), and mtDNA (dotted circle in Figure S21, dotted circle and arrow in Figure S22, haplotypes within L0d1c haplogroup indicated with arrows 2-6 in Figure S23, and haplotypes within L0d2a1 in Figure S24). These findings are in agreement with analyses based on autosomal data indicating that the Naro are genetically intermediate between northwestern Kalahari (i.e. Ju|’hoan and !Xuun) and southeastern Kalahari (Taa) populations, just as they are intermediate geographically (Pickrell et al., 2012). Even though they show high genetic affinities to the Kx’a and Tuu-speaking populations, Naro speak a Khoe-Kwadi language. The Naro kinship system has been described as a simplified Khoe kinship system with some Ju|’hoan features, and it has been hypothesized that they may have spoken a Kx’a language in the past and subsequently shifted to a Khoe-Kwadi language (Güldemann, 2008; Barnard, 2016). Genetic data indicate that this culture and language shift was not just a cultural process, but rather that it was driven by the diffusion of the Khoe-Kwadi-speaking people. Possible genetic evidence of contact between proto-Naro, who may have spoken a Kx’a-related language, with Khoe-Kwadi populations that could have contributed to the language shift in Naro may be found in branch 11 of the network for NRY haplogroup A3b1 (Figure S4). This branch harbors almost exclusively haplotypes from current Khoe-Kwadi speakers (i.e. Naro, ║Ani/Buga, and ║Xo, who all belong to the West Kalahari Khoe sub-branch; Figure S1). Although most mtDNA lineages in the Naro are shared with Kx’a- and Tuu-speaking groups, there are some lineages that are shared with Khoe-Kwadi-speaking groups, i.e. in one L0k lineage shared with Khwe and G║ana (arrow in Figure S21), one L0d1c haplotype predominantly shared with G|ui and East Kalahari Khoe speakers (arrow 1 in Figure S23), and for L0d2ab there are haplotypes shared or in close proximity to haplotypes from Hai║om and G|ui (Figure S24). Both the mtDNA and the Y-chromosome evidence thus suggests that the putative language shift in the Naro was accompanied by some gene flow.

#### G║ana and Kgalagadi

The G║ana are multilingual, like most Khoisan populations; they speak both their own language (in this case, a Khoe-Kwadi language) and a Bantu language (in this case, Kgalagadi). Based on their mtDNA haplogroup composition and Φst values they are similar to other Khoisan populations, but they are a clear outlier for the NRY (Figure 1, Figure 4 and Figure S11, Table S11). The proximity of the G║ana to the Kgalagadi in the NRY-based NJ tree and CA (Figure S11 B and D) is in agreement with their own belief that the intermarriage between Khoisan females (presumably G|ui or their close relatives) and Bantu males (presumably Kgalagadi) resulted in the founding of the G║ana population (Barnard, 1992). This cultural belief of extreme sex-biased admixture is supported by the high frequency of NRY haplogroup B2a in G║ana (80%) and the fact that the lowest pairwise Φst values for the NRY between G║ana and any other population is with the Bantu-speaking Kgalagadi (Table S11). The relatively low mtDNA Φst values between G║ana and other Khoisan groups and correspondingly closer relationships (Table 1, Figure S11 A and C) are consistent with their having Khoisan matrilineal ancestry. This mixed ancestry for the G║ana is also reflected in the autosomal data, as they have an estimated 30-41% Bantu-related ancestry (Pickrell et al., 2014; Uren et al., 2016). As they have exclusively Khoisan-related mtDNA lineages L0d and L0k (Barbieri et al., 2013b, 2014), the probable source of autosomal Bantu ancestry is via male lineages. Since clear Bantu-related NRY lineages are in lower frequency than the estimated Bantu-related autosomal ancestry, this raises the possibility that at least some of the B2a lineages entered the G║ana population from surrounding Bantu populations. If all or most of the B2a haplotypes found in G║ana came from Bantu populations, then the G║ana would represent the population that experienced the most extreme case of sex-biased gene flow (see Figure 4 with B2a as NAUT).

The Kgalagadi show the highest proportion of autochthonous uniparental haplogroups among Bantu populations (Figure 4). Interestingly, even though some authors argue that the direct ancestors of the Tswana and Kgalagadi probably migrated to what is now Botswana as recently as 350 years ago (Kiyaga-Mulindwa, 1993; Segobye, 1998), Barnard (1992) has suggested that the Kgalagadi are the oldest existing Bantu-speaking inhabitants of Botswana and entered the southern part of the country probably centuries before European colonization. If this scenario is correct, the putative long period of cohabitation between Kgalagadi and local foraging groups in the area could explain the relatively high proportion of autochthonous uniparental haplogroups found in the Kgalagadi.

### Multiple waves of sex-biased gene flow in the context of the putative East African pastoralist migration and the Bantu expansion

Southern African populations exhibit a complex genetic makeup characterized by strong sex-biased gene flow (Figure 4 and Figure 5), reflecting a long history of population movements and interactions between local and incoming populations. There are at least two migration events that could have contributed to the complex and sex-biased ancestry of Khoisan populations: the Bantu expansion; and an earlier migration from eastern Africa that is putatively associated with the introduction of pastoralism (Pickrell et al., 2014).

#### Admixture with East African migrants

Archaeological and linguistic evidence has suggested a pre-Bantu migration from eastern Africa that brought pastoralism and Khoe-Kwadi languages (Mitchell, 2002; Blench, 2006; Güldemann, 2008; Pleurdeau et al., 2012), and thus had a significant impact on southern Africa. Although studies of different genetic markers support the demic migration model from East Africa, they vary in their conclusions concerning the impact and importance of this migration (see Introduction). For instance, the mtDNA data support very limited gene flow (Barbieri et al., 2014; Uren et al., 2016), while the lactase persistence and limited NRY data undeniably support the demic diffusion model with significant population movement (Henn et al., 2008; Schlebusch et al., 2012; Breton et al., 2014; Macholdt et al., 2014, 2015). Genome-wide data from both ancient remains and modern populations support the demic diffusion model, but with various interpretations of its significance (Schlebusch et al., 2012, 2017; Uren et al., 2016; Montinaro et al., 2017; Skoglund et al., 2017).

One possibility is that the signal of East African ancestry in Khoisan populations was shaped by a heavily male-mediated migration from eastern Africa (as previously proposed in Barbieri et al., 2014). Thus, it is crucial to investigate the paternal history of Khoisan populations in order to differentiate between the spread of pastoralism due to limited demic migration with more significant cultural diffusion vs. a heavily male-biased demic migration from the east that brought pastoralism. We do not find a higher frequency of the E1b1b NRY haplogroup, previously associated with the spread of pastoralism (Henn et al., 2008), in the pastoralist Nama (Table S8, Table S9). Even though the Nama are the only sensu stricto Khoe-speaking pastoralist population, they harbor just a subset of the E1b1b NRY diversity compared with other Khoisan populations (Figure 3, Figure S10). This is contrary to the expectation that diversity should be highest in the population that most probably brought pastoralism, or represents a direct descendant of such a population. If E1b1b was indeed associated with a migration of pastoralists, subsequent demographic events and/or changes in subsistence practices in southern Africa have diminished the association.

However, even though it cannot be directly associated with pastoralism, haplogroup E1b1b clearly has an eastern African origin. The close relationship of southern African E1b1b STR haplotypes to haplotypes from the two eastern African foraging populations, Hadza and Sandawe (Figure 3), indicate a common origin of haplotypes in southern and eastern Africa. This is in agreement with the linguistic hypothesis of a relationship between proto-Khoe-Kwadi and Sandawe (Güldemann and Elderkin, 2010). Interestingly, the diversity of haplotypes seen in different Khoisan populations, with multiple star-like expansions from haplotypes in close proximity to eastern African foragers, suggest a more complex migration history for haplogroup E1b1b than previously suspected. This migration may have included multiple distantly-related haplotypes that subsequently were sorted into different populations, and/or there may have been more than one migration event connecting eastern with southern Africa. Further studies are needed to clarify the relationships between eastern and southern African populations.

#### Eurasian haplogroups

The Nama are unusual in having the highest frequency of Eurasian NRY haplogroups (32%); the Damara (who live in close association with the Nama) have the next highest frequency (14%), and no other group has more than one Eurasian NRY sequence (Table S6). The high frequency of Eurasian haplogroups in the Nama is in agreement with autosomal data that show they have a relatively high Eurasian ancestry component (Pickrell et al., 2012, 2014; Uren et al., 2016; Montinaro et al., 2017). These findings could be explained by recent admixture with colonialists and/or with an already admixed population from eastern Africa (Wallace and Kinahan, 2011; Pickrell et al., 2014). Even though some of the Y chromosome haplogroups were most likely brought by European colonists (e.g. haplogroups O, G2a2b2a1, I1, I2a2a, R1a; Underhill and Kivisild, 2007; Capredon et al., 2013) others could also represent traces of migration from elsewhere in Africa (e.g. R1b1a2, and/or T1; Karafet et al., 2008; Chiaroni et al., 2009; Cruciani et al., 2010; Gomes et al., 2010; Mendez et al., 2011; Capredon et al., 2013).

#### Bantu Expansion

Our results confirm and extend previous findings concerning the enormous impact that the spread of iron-using agro-pastoralist populations speaking Bantu languages had on the genetic landscape of southern Africa (Beleza et al., 2005; Wood et al., 2005; Coelho et al., 2009; de Filippo et al., 2011; Schlebusch et al., 2011, 2017, Pickrell et al., 2012, 2014; Barbieri et al., 2014; Marks et al., 2014). Arriving in southern Africa ~2,000-1,200 years ago (Reid et al., 1998; Phillipson, 2005; Kinahan, 2011), the Bantu expansion resulted in a sex-biased pattern of admixture that is characterized by a high proportion of autochthonous mtDNA lineages in agro-pastoralist populations, and a high proportion of non-autochthonous NRY lineages in foragers (Figure 4). However, the intensity of sex-biased admixture varies widely among populations (Figure 4) and moreover shows geographic structure (Figure 5 and Figure S12), indicating that this was a complex process.

Numerous differences in cultural and sex-specific practices could have contributed to shaping the current genetic pattern seen in expanding agriculturalist Bantu-speaking populations and local forager populations (reviewed in Heyer et al., 2012). Based on comparisons of pairwise Φ_ST_ distances for the mtDNA and NRY and calculations of effective migration rates between Bantu and Khoisan populations (ν_mtDNA_/ν_NRY_), our results suggest that there has been more male-biased migration from Bantu to Khoisan populations than female-biased migration from Khoisan to Bantu populations. It should be noted that the estimates of effective migration rates should be viewed with caution, as the model underlying this approach is based on many biologically unrealistic assumptions (Whitlock and McCauley, 1999; Holsinger and Weir, 2009) and because the method relies on F_ST_ (or Φ_ST_) values that can also be used to estimate coalescence times between populations (Slatkin, 1991). Thus, F_ST_ alone cannot be used to conclude whether the relatedness of populations is due to ongoing migration, recent common ancestry, or both (reviewed in Holsinger and Weir, 2009). Nevertheless, previous autosomal studies, as well as the presence of non-autochthonous uniparental lineages in autochthonous populations and vice versa, testify to the impact of gene flow between populations, and thus it is very likely that differences in Φ_ST_ values are largely influenced by migration. Therefore, it is reasonable to assume that the calculated Φ_ST_ values at least partly reflect the higher rate of male-biased than female-biased migration between Khoisan and Bantu populations. Females from forager communities are known to be preferred by Bantu-speaking males because of their reputation for greater fertility, and the lower (if any) bride price of forager wives (Cavalli-Sforza, 1986; Lee, 1993; Destro-Bisol et al., 2004). The Bantu to forager flow of paternal lineages occurs if the children of such liaisons remain in the forager villages (Lee, 1993). Conversely, the flow of maternal lineages from forager to Bantu groups occurs if the children are brought up in the Bantu communities. Strong sociocultural taboos inhibit unions between forager males and Bantu females (Cavalli-Sforza, 1986; Lee, 1993). Although this expected sex-biased signature is stronger in Khoisan than in Bantu populations, there is considerable variation in the intensity of the sex bias among different populations (Figure 5C, Figure S12C), and so other factors must also play a role.

Residential practices is another cultural trait that is likely to influence the distribution of genetic variation, and thus the signal of sex-biased gene flow. Patrilocal populations are expected to show less population differentiation for the mtDNA than the NRY because of higher rates of female migration between local groups, while the reverse is expected for matrilocal populations. Indeed, it has been observed that mtDNA is more geographically structured than the NRY in matrilocal populations (Oota et al., 2001; Bolnick et al., 2006; Kumar et al., 2006), while patrilocal populations show contrasting patterns (Wilder et al., 2004; Kumar et al., 2006; Langergraber et al., 2007). Even though our data set consists of patrilocal Bantu populations and Khoisan populations that preferentially practice patrilocal postmarital residence after an initial period of matrilocality (and to a lesser extent neolocality; Barnard, 1992), we observe larger differences among populations for mtDNA than for the NRY (Table 1), and thus higher effective migration rates for NRY than for mtDNA (Table S10), which is the pattern characteristic for matrilocality and not for patrilocality. Even more intriguingly, we observe that the putatively higher NRY than mtDNA migration is more prominent between Bantu populations than between Khoisan populations. This deviation could be explained by a rapid male-dominated Bantu expansion over huge geographic areas and incorporation of already geographically structured mtDNA lineages into expanding Bantu populations. This scenario would result in a relatively homogeneous NRY genetic pool in Bantu populations across a huge geographic area, yet with more diverse and geographically structured mtDNA. Marks et al., (2012) showed that even though female migration is more frequent among patrilocal populations, males migrate preferentially at longer distances than females, suggesting that patrilocal residence is expected to mostly impact geographically close groups. As geographic distances between populations in our data set are mostly >200km, which would be considered long-range distances (Marks et al., 2012), it is possible that the observed pattern is due to a higher migration rate of men at longer geographic distances rather than an overall higher migration rate. However, our data are insufficient for separating the effects of migration rate vs. geographic distance; further studies are needed.

One of the most striking findings of our study is the increase of sex-biased admixture from north to south. With the assumption that the initial contact between Bantu and Khoisan populations occurred in the north, the increasing ISBGF towards the south (Figure 5 A-B, Figure S12 A-B) could be interpreted as an indication that the initial contact involved less sex bias. The north to south increase in autochthonous uniparental lineages in Bantu populations (Figure S16) could suggest either more intense intermarriage with local populations in the south, and/or a gradual accumulation of autochthonous uniparental markers in Bantu populations towards the south as a result of interactions between the southwards migrating Bantu populations and autochthonous populations that they encountered on their way. The former seems more likely, as autochthonous lineages incorporated in Bantu populations tend to be regionally specific, e.g. L0k is found mainly in the north, both in Khoisan and Bantu groups, but not in the south Bantu groups (Barbieri et al., 2013b; Schlebusch et al., 2013), in contrast to what would be expected if autochthonous lineages had gradually accumulated during the southward migration of Bantu-speaking groups. This is compatible with the Static and Moving frontiers model that implies limited assimilation of foragers into agro-pastoralist populations during the moving frontier phase, and increased likelihood of gene flow between populations with the occurrence of a static frontier (Marks et al., 2014). It is also consistent with Destro-Bisol et al’s (2004) model that proposes the gradual establishment of social inequalities between agro-pastoralists and foragers that resulted in asymmetric gene flow between them. During the early phase of contact between agro-pastoralists and foragers, the survival of the newly arrived food producing societies in an unfamiliar habitat was probably heavily dependent on the knowledge of autochthonous foragers, and as such their contact might have been more egalitarian. It has been suggested that Bantu languages of the Kavango-Zambezi transfrontier region that have incorporated click consonants might have acquired them during this more egalitarian phase, when Khoisan populations had higher social status (Pakendorf et al., 2017). Later, as Bantu groups became more established and less dependent on local knowledge, they became more socially dominant, and hence sex bias increased in intensity due to the establishment of socio-cultural taboos (Destro-Bisol et al., 2004). Thus, under this interpretation the geographic pattern reflects an increase in sex bias over time.

In conclusion, we have carried out a comprehensive population-level study of matrilineal and patrilineal lineages in southern African populations, integrated within their historical and anthropological background. Discrepancies found between the linguistic and genetic relationships of Damara, Naro, ǂHoan, and Hai║om suggest probable language shift and/or extensive contact between these and other, linguistically unrelated, populations. We find support for a migration from eastern Africa but do not find an association of NRY haplogroup E1b1b with pastoralism today, suggesting that the arrival of pastoralism was more complex than previously suspected. Our study indicates that the Bantu expansion was probably a rapid, male-dominated expansion, during which local Khoisan females were much more likely to be absorbed into Bantu populations than Khoisan males. We detected a stronger intensity of sex-biased gene flow in Khoisan populations than in Bantu populations via the incorporation of non-autochthonous NRY lineages into Khoisan populations. Finally, we find that the intensity of the sex-biased gene flow increases from north to south, possibly due to the more recent arrival of Bantu populations in the south along with the gradual establishment of social inequalities between autochthonous Khoisan populations and expanding Bantu-speaking populations. Further studies with ancient samples spanning the time frame from pre-Eastern African migration to post-Bantu migration, along with further analyses of modern samples, will clarify the temporal dynamics of interactions between these migrations and the autochthonous populations.

## Acknowledgements

This study focuses on the genetic prehistory of populations and it does not intend to evaluate or influence the cultural identity of any group, which consists of much more than just genetic ancestry. We thank all sample donors for their invaluable contribution, and members of the Human Population History Group for fruitful discussions and comments. This work was carried out within the EUROCORES Programme EuroBABEL of the European Science Foundation and was supported by funds from the Deutsche Forschungsgemeinschaft and the Max Planck Society. We furthermore thank the Wenner-Gren Foundation and the LABEX ASLAN for supporting a return trip to Botswana and Namibia to share our results with the communities who participated in the study. BP is grateful to the LABEX ASLAN (ANR-10-LABX-0081) of Université de Lyon for its financial support within the program “Investissements d’Avenir” (ANR-11-IDEX-0007) of the French government operated by the National Research Agency (ANR).

